# An ART-fold Rhs toxin from *Pluralibacter gergoviae* defines Tne5, a new clade of NAD(P)⁺ glycohydrolases effectors

**DOI:** 10.64898/2026.01.26.701734

**Authors:** Jonas B. Desjardins, Martin Durrmeyer, Eric Cascales

## Abstract

The type VI secretion system (T6SS) ia a widespread bacterial nanomachine that mediates interbacterial competition by delivering toxic effectors into neighboring cells. Among these, enzymes targeting nicotinamide adenine dinucleotide cofactors (NAD⁺ and NADP⁺) are particularly potent because they rapidly disrupt redox homeostasis and central metabolism. Several families of T6SS-associated NAD(P)⁺-consuming effectors (Tne1-Tne4) have been described. Here, we characterize a T6SS-associated Rhs toxin from *Pluralibacter gergoviae*. Competition assays show that *P. gergoviae* kills *Escherichia coli* in a T6SS-dependent manner. Heterologous production reveals that the Rhs C-terminal extension is toxic in the *E. coli* cytoplasm and that co-production with the protein encoded downstream neutralizes this activity. AlphaFold3 modeling predicts that the toxin adopts an ADP-ribosyltransferase (ART)-like α/β fold with a putative catalytic pocket accommodating NAD. By contrast to T6SS ART toxins described so far, the toxin does not inhibit transcription, translation or cell division, but instead depletes NAD⁺ and NADP⁺. Phylogenetic analyses and structural modeling show that this effector defines a new ART-related family of NAD(P)⁺ glycohydrolases, which we propose to name Tne5, broadly distributed across antagonistic systems.

## INTRODUCTION

Bacteria deploy antibacterial weapons to compete for nutrients and space in densely populated niches (1–5). One of the key actors of bacterial competition is the type VI secretion system (T6SS), a sophisticated nanomachine belonging to the broad family of contractile injection systems, which uses a contractile mechanism to deliver effectors into neighboring cells (6–9). The activity of the T6SS impacts the composition and organisation of microbial communities, provides a privileged access to nutrients, and promotes colonization of the niche (4, 10–13). The T6SS is comprised of a membrane complex that anchors a cytoplasmic phage-related structure made of an assembly platform, the baseplate, and of the tail tube/sheath complex (6–8, 14). The tail tube/sheath complex is composed of a needle wrapped by the contractile sheath, whose contraction propels the needle towards target cells (7, 15). The needle is made of an inner tube of stacked Hcp hexamers tipped by a VgrG-PAAR spike complex (7, 8, 15). The effectors are either independent polypeptides that associates, directly or through the help of adaptors and chaperones, to the needle, or domain fused to structural subunits of the needle, Hcp, VgrG or PAAR (9, 16–19). These later effectors belong to the family of polymorphic toxins (PTs), which are comprised of a C-terminal effector domain fused to an N-terminal trafficking domain that target the toxin to the proper delivery apparatus (20). Beyond these trafficking and toxic domains, PTs may contain supplementary structural elements. For example, T6SS-associated PAAR PTs commonly possess transmembrane helices that are proposed to promote insertion into target cell membranes, or Rearrangement hot spot (Rhs) domains that form β-barrel cocoons encapsulating the toxic C-terminal extension (21–27).

Antibacterial T6SS effectors target essential molecules, macromolecules and processes of the bacterial physiology: cell wall integrity, membrane homeostasis, nucleic acids, cofactors and nucleotides, or the translation machinery (9, 19, 28–30). Within this diversity, a broad family of T6SS enzymatic effectors are related to nicotinamide adenine dinucleotide (NAD⁺) or NAD+ phosphate (NADP⁺) (31). This family includes ADP-ribosyltransferases (ART) that catalyze the transfer of ADP-ribose from NAD⁺ to proteins or nucleic acids (32–41). Structurally, ARTs fold as a split β-sheet typically composed of six strands in the order β4-β5-β2 / β1-β3-β6, presenting diverse catalytic triads (H-H-h, H-Y-[QED], R-[ST] or R-H-h) that define major lineages (32, 33, 36). In addition, other highly potent effectors with ART-like folds can catalyze NAD^+^ or NADP^+^ degradation, such as NAD(P)⁺ glycohydrolases (NADases), or cyclization reactions, such as ADP-ribosyl cyclases (ARC) (42–47). These NADase or ARC activities rapidly compromise redox balance and metabolism by depleting NAD(P)^+^ at high rates, leading to growth arrest or death (31). Recent comparative analyses have classified T6SS-associated NAD(P)⁺ glycohydrolases into four major families (<u>T</u>ype VI secretion <u>N</u>ADase <u>e</u>ffectors 1-4, Tne1-Tne4) based on phylogenetic analyses (42, 43, 45).

Here, we show that *Pluralibacter gergoviae* ATCC33028 (formerly *Enterobacter gergoviae*) carries a T6SS gene cluster encoding a putative polymorphic toxin comprising PAAR and Rhs domains, Rhs^Pg^. Competition assays demonstrate that the *P. gergoviae* T6SS is active and eliminates *E. coli*. Heterologous expression in *E. coli* shows that the Rhs^Pg^ C-terminal extension (Rhs^Pg^-CT) is toxic and can be neutralized by the protein encoded downstream of *rhs*. Rhs^Pg^-CT AlphaFold3 structural modeling predicts that the domain adopts an ART-like α/β fold with a putative catalytic cleft accommodating NAD, comprising residues Lys1392, and Glu1482. Mutational analysis further shows that Lys1392 is important for toxicity, while Glu1482 is not. Functional assays demonstrate that Rhs^Pg^-CT does not inhibit transcription, translation or cell division and does not ADP-ribosylate target cell proteins, but rather depletes NAD⁺ and NADP⁺ *in vitro*, classifying it as a member of the Tne family. Phylogenetic analyses and structural comparisons further show this Tne is evolutionarily closer to ART enzymes than previously described Tne families, defining a new family of NAD(P)⁺ glycohydrolases, which we term Tne5, and shedding light on the diversification of ART-like folds in bacterial competition.

## RESULTS

### A T6SS locus in *Pluralibacter gergoviae* mediates antibacterial activity

*Pluralibacter gergoviae* is an environmental Enterobacterales species that can occasionally cause opportunistic urinary tract infections (48). Because of its high tolerance to widely used preservatives such as parabens and isothiazolinones (49–51), *P. gergoviae* has been associated with multiple contamination outbreaks and recalls of cosmetic, personal care and hygiene products (52–54). Genome screening of *P. gergoviae* ATCC 33028 revealed a T6SS gene cluster containing the core genes (*tssA-M*), accessory modules (*tag* genes), and toxin-immunity pairs such as a putative Tae4 amidase, consistent with a functional antibacterial system (Fig. 1*A*). Indeed, CPRG reporter-based antibacterial competition assays demonstrated that wild-type (WT) *P. gergoviae* eliminated *Escherichia coli* K12 W3110 whereas its isogenic *tssL* mutant did not (Fig. 1*B*), demonstrating T6SS-dependent antibacterial activity.

**Figure 1.**
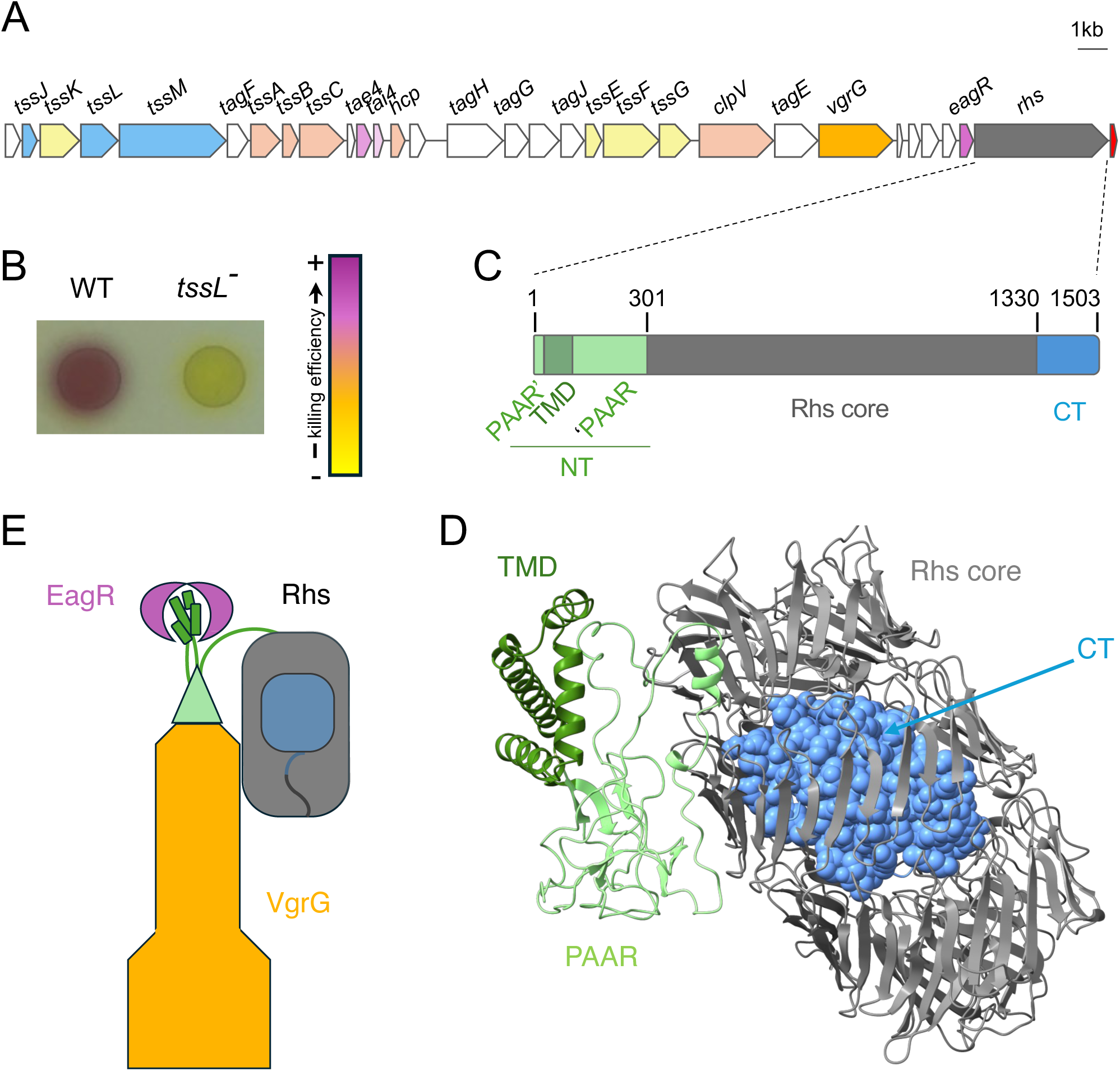
A T6SS locus in *Pluralibacter gergoviae* encodes a putative Rhs toxin-immunity pair and mediates antibacterial activity. **(A)** Schematic of the *P. gergoviae* ATCC 33028 T6SS gene cluster. The different *tss* (blue, membrane complex; yellow, baseplate; flesh, tail tube/sheath) and *tag* (white) genes are indicated, as well the genes encoding a VgrG spike (orange), a predicted Tae4-Tai4 pair (pink), a EagR chaperone (purple), a Rhs protein (grey) and its putative immunity (red). **(B)** LAGA interbacterial competition assay. Wild-type (WT) *P. gergoviae* or its isogenic *tssL* mutant were mixed with LacZ⁺ *E. coli* recipients, spotted on LB agar plates and incubated for 4 hours at 37°C. Recipient cell lysis after competition was revealed using CPRG (yellow), which turns purple upon hydrolysis by β-galactosidase from lysed *lacZ^+^* recipient cells. The killing efficiency heatmap (from yellow (no killing) to purple (killing)) is shown on the right. **(C and D)** Schematic representation (**C**) and AlphaFold3 structural model (**D**) of the Rhs^Pg^ protein, highlighting the different domains and their boundaries: PAAR domain (PAAR’ or prePAAR and ‘PAAR in light green), trans-membrane domain (TMD in dark green), Rhs core (grey), and C-terminal extension (blue). The confidence scores, pLDDT and PAE are presented in Supplemental Fig. S1. **(E)** Schematic representation of the Rhs^Pg^ effector (same colors as panels C and D) loaded to the VgrG spike (orange). A dimer of EagR chaperones (purple) is shown bound to the Rhs^Pg^ TMD. The AlphaFold3 model of the VgrG-Rhs-EagR complex, and its confidence scores, pLDDT and PAE are presented in Supplemental Fig. S2.

### The *P. gergoviae* T6SS encodes a Rhs polymorphic toxin

The *P. gergoviae* T6SS gene cluster carries a *rhs* gene (Rhs^Pg^, NCBI Reference Sequence: WP_330993187; GeneBank Identifier (GI): AVR02352; locus tag: A8H26_06345) immediately followed by a small gene encoding a predicted cytoplasmic protein (GI: AVR02353; locus tag: A8H26_06350), a gene organization typical of Rhs-immunity pairs. Domain annotation of the 1503-amino-acid (aa) Rhs^Pg^ protein demonstrated that it shares the canonical modular organization of T6SS Rhs toxins (24–27): (i) an N-terminal region (amino-acids (aa) 1-301) comprising a PAAR domain (PAAR’, aa 1-15 and ‘PAAR, aa 98-301) that likely specifies transport by the T6SS through association with the VgrG spike protein (55–57) and a predicted 4-helix transmembrane domain (TMD, aa 16-97), in agreement with the presence of a gene upstream *rhs* encoding a putative TMD-protecting chaperone EagR (21, 22), (ii) a central YD-repeat Rhs encapsulation β-barrel (aa 302-1329), and (iii) a C-terminal extension (aa 1330-1503, Rhs^Pg^-CT) that typically carries the toxic activity (Fig. 1, *C* and *D*). These three regions are separated by two cleavage sites that are responsible for the release of the buried C-terminal extension: the N-terminal PVHAATGV motif resembling the PVSMVTGE and PVYVASGE signatures of *Aeromonas dhakensis* and *Photorhabdus laumondii* Rhs proteins (24, 58), and the canonical autoproteolysis DPVGL-X_17_-DPWGL C-terminal motif (24, 58). AlphaFold3 modeling prediction confirmed the canonical structural organization of Rhs proteins, with the C-terminal extension encapsulated within the Rhs core β-barrel (Fig. 1*D* and Fig. S1). This prediction also suggested that the N-terminal domain has a typical PAAR fold, that potentially binds to the tip of the VgrG protein encoded 6 genes upstream of *rhs* (Fig. S2). Finally, as experimentally demonstrated for the *Pseudomonas aeruginosa* Tse6 effector (21, 22), two EagR subunits form a horseshoe-shaped protecting shell around the predicted TMD (Fig. S2). Overall, this organisation suggests that the Rhs polymorphic toxin uses its PAAR domain to bind to VgrG for its translocation, with the EagR chaperones required for maintaining the TMD in a state competent for translocation (Fig. 1*E*).

To test whether the Rhs^Pg^ C-terminal extension has antibacterial activity, the sequence downstream the C-terminal autoproteolysis motif (corresponding to aa 1330-1503) was cloned into the low-copy, IPTG-inducible, pNDM220 vector and heterologously produced in the *E. coli* cytoplasm. Figure 2 shows that induction of the expression of Rhs^Pg^-CT was toxic in *E. coli*. Co-expression of the A8H26_06350 gene, located downstream of *rhs*, from the pBAD33 vector, restored growth (Fig. 2), suggesting it neutralizes Rhs^Pg^-CT toxicity. Taken together, these results demonstrate that Rhs^Pg^-CT is an antibacterial toxin acting in the cytoplasm, whereas A8H26_06350 encodes its cognate immunity protein.

**Figure 2.**
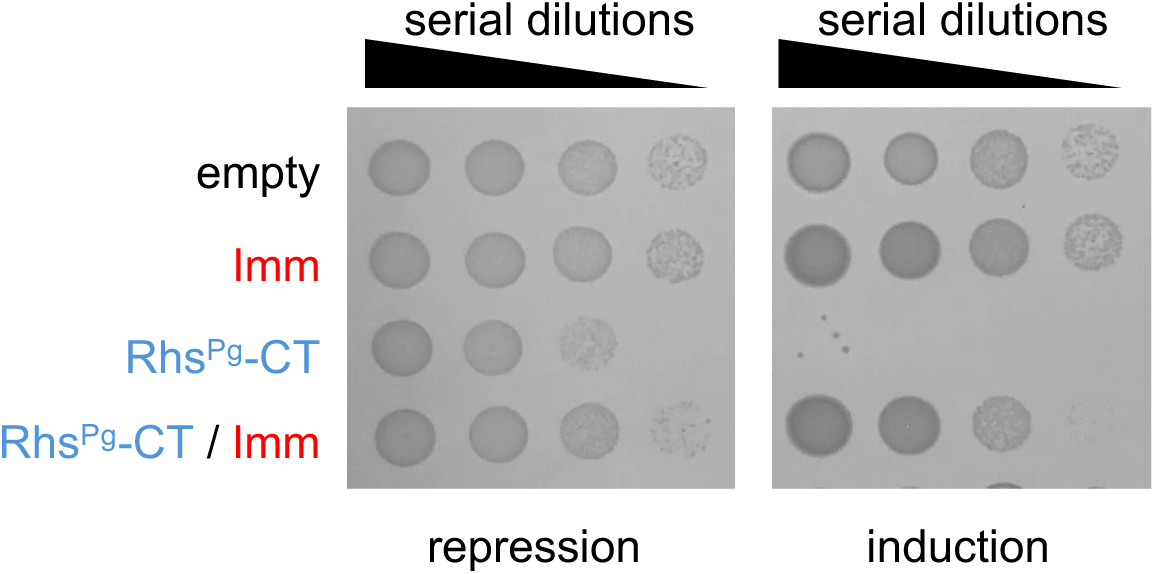
The Rhs^Pg^ C-terminal extension is toxic in the *E. coli* cytoplasm and is neutralized by the downstream immunity protein. Toxicity assay in the heterologous host *E. coli.* Overnight cultures of *E. coli* cells producing the Rhs^Pg^ C-terminal extension (Rhs^Pg^-CT) from the low-copy vector pNDM220, the putative immunity protein (A8H26_06350, Imm) from the pBAD33 vector, or both were serially diluted (10^-1^ to 10^-4^) and spotted on LB agar plates supplemented with 1% of glucose (left panel, repression), or with 0.05 mM of IPTG and 0.2 % of L-arabinose (right panel, induction). Empty, empty pNDM220 and pBAD33 vectors.

### Rhs^Pg^-CT shares a typical ADP-ribosyltransferase fold

AlphaFold3 modeling of Rhs^Pg^-CT revealed an ADP-ribosyltransferase (ART)-like fold, comprising a compact α/β catalytic scaffold built around a split β-sheet delimiting a cleft and flanked by 5 α-helices (Fig. 3*A*). As in typical ARTs, the β-sheet core is composed of six β-strands organized as two perpendicular 3-stranded β-sheets in the order β4-β5-β2 / β1-β3-β6 (Fig. 3*A*). However, β-strand β4 is missing or not predicted by AlphaFold3 in Rhs^Pg^-CT (Fig. 3*A*). In ART enzymes, NAD⁺ positioning and catalysis rely on a triad of residues usually located on strands β1, β2 and β5, defining major families (H-Y-E, R-S-E, and h-h-H) (32). Sequence analyses of Rhs^Pg^-CT suggested that Lys1392 and Glu1482 may constitute the putative catalytic pocket (Fig. 3*B*). A highly confident model including NAD⁺ (ipTM=0.96) positionned the dinucleotide in the Rhs^Pg^-CT cleft (Fig. 3*C*) whereas the AlphaFold3 co-model of Rhs^Pg^-CT with its immunity protein (ipTM=0.94) predicted formation of a tight complex between the two partners, in which a loop of the immunity penetrates deeply into the Rhs^Pg^-CT cleft, sterically occluding the putative NAD^+^-binding pocket (Fig. 3*D*). Alanine substitution of Rhs^Pg^-CT residue Glu1482 had a very weak reduction in activity, whereas the K1392A mutation significantly impaired toxicity (Fig. 3*E*).

**Figure 3.**
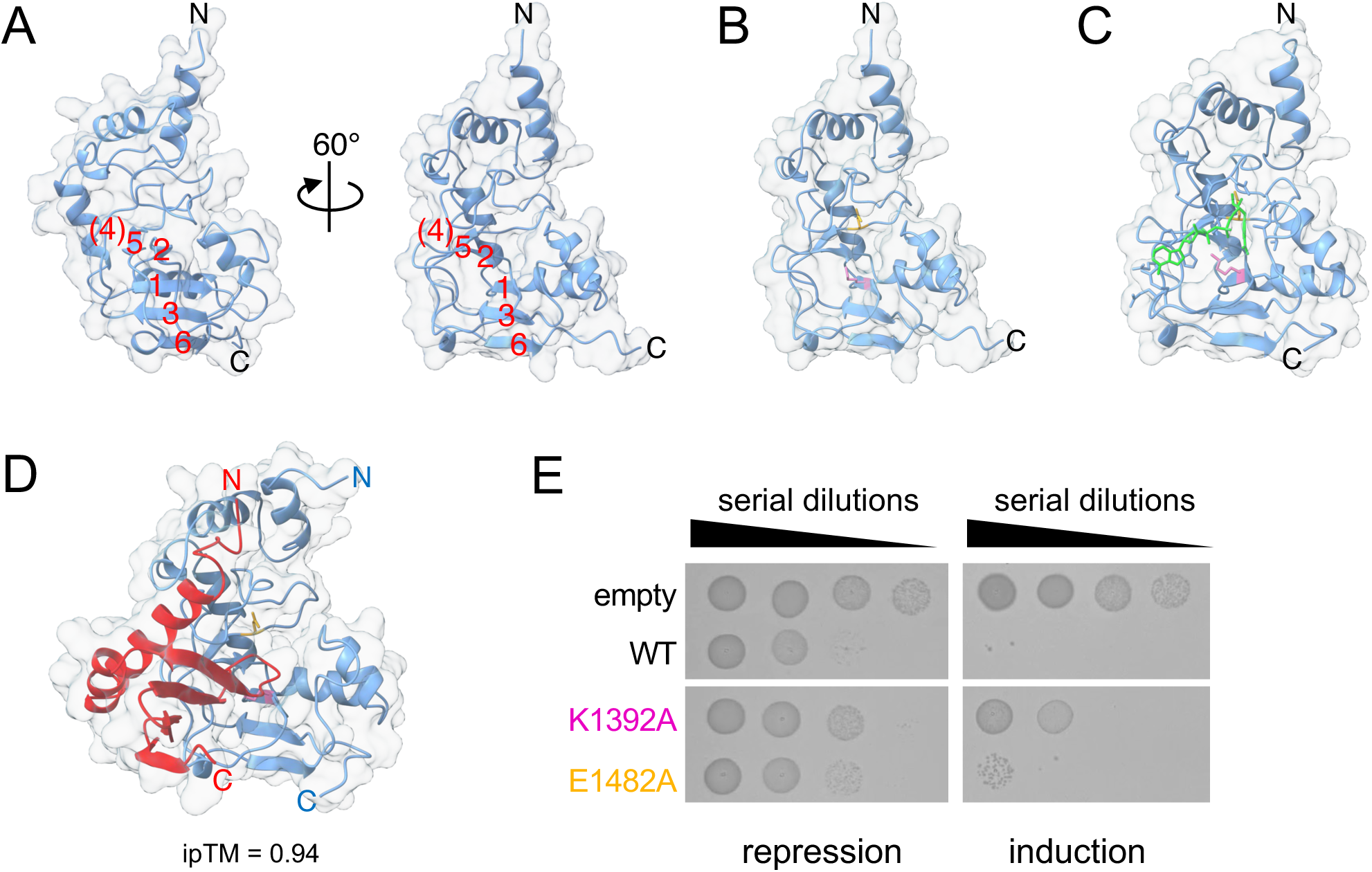
RhsPg-CT shares an ART fold with a putative catalytic site accommodating the NAD^+^ and occluded by its cognate immunity. (**A**) Ribbon representation of the AlphaFold3 prediction model of Rhs^Pg^-CT. The β-strands of the split β-sheet are numbered in red. The predicted region located at the position of β-strand β4 in the typical ART fold is indicated in brackets. The surface structure is shown in grey transparent. N, N-terminus; C, C-terminus. The confidence score, pLDDT and PAE are presented in Supplemental Fig. 3A-B. (**B**) Ribbon representation of the AlphaFold3 prediction model of Rhs^Pg^-CT highlighting the positions of residues K1392 (pink) and E1482 (orange). (**C**) Ribbon representation of the AlphaFold3 prediction model of Rhs^Pg^-CT bound to NADH (ipTM score = 0.96). The Rhs^Pg^-CT residues K1392 and E1482 are shown in pink and orange, whereas the NADH is shown in green. The confidence score, pLDDT and PAE are presented in Supplemental Fig. 3C-D. **(D)** Ribbon representations of the AlphaFold3 prediction model of the Rhs^Pg^-CT/Imm complex (ipTM score = 0.94). The toxin domain is shown in blue whereas the immunity is shown in red. The surface structure is shown in grey transparent. N, N-terminus; C, C-terminus. The confidence score, pLDDT and PAE are presented in Supplemental Fig. 3E-F. (**E**) Toxicity assay in the heterologous host *E. coli.* Overnight cultures of *E. coli* cells producing wild-type (WT) Rhs^Pg^-CT or the K1392A or E1482A variants from the low-copy vector pNDM220 were serially diluted (10^-1^ to 10^-4^) and spotted on LB agar plates, supplemented with 1% of glucose (left panel, repression), or with 0.05 mM of IPTG (right panel, induction). Empty, empty pNDM220 vector.

### Rhs^Pg^-CT does not inhibit transcription, translation or cell division

Given that most T6SS-associated ART-fold toxins have been shown to inhibit translation in a NAD^+^-dependent manner, through the modification of RNAs or translation factors (38–40), we first asked whether Rhs^Pg^-CT impacts protein synthesis using a coupled *in vitro* transcription-translation (IVT) assay. Rhs^Pg^-CT was first synthesized by IVT in the absence of NAD^+^ and then incubated in a new IVT reaction in the presence of a DNA template encoding GFP. Figure 4*A* shows that Rhs^Pg^-CT did not prevent GFP production in the presence or absence of NAD^+^, demonstrating that Rhs^Pg^-CT does not inhibit transcription and translation. To identify potential protein targets of Rhs^Pg^-CT, we incubated IVT-synthesized Rhs^Pg^-CT with *E. coli* cell lysates in the presence of 6-biotin-17-NAD⁺, a NAD^+^ derivative in which the ADP-ribose moiety is decorated with a biotin. The biotinylated protein profiles did not differ detectably in the absence or presence of Rhs^Pg^-CT (Fig. 4*B*).

**Figure 4.**
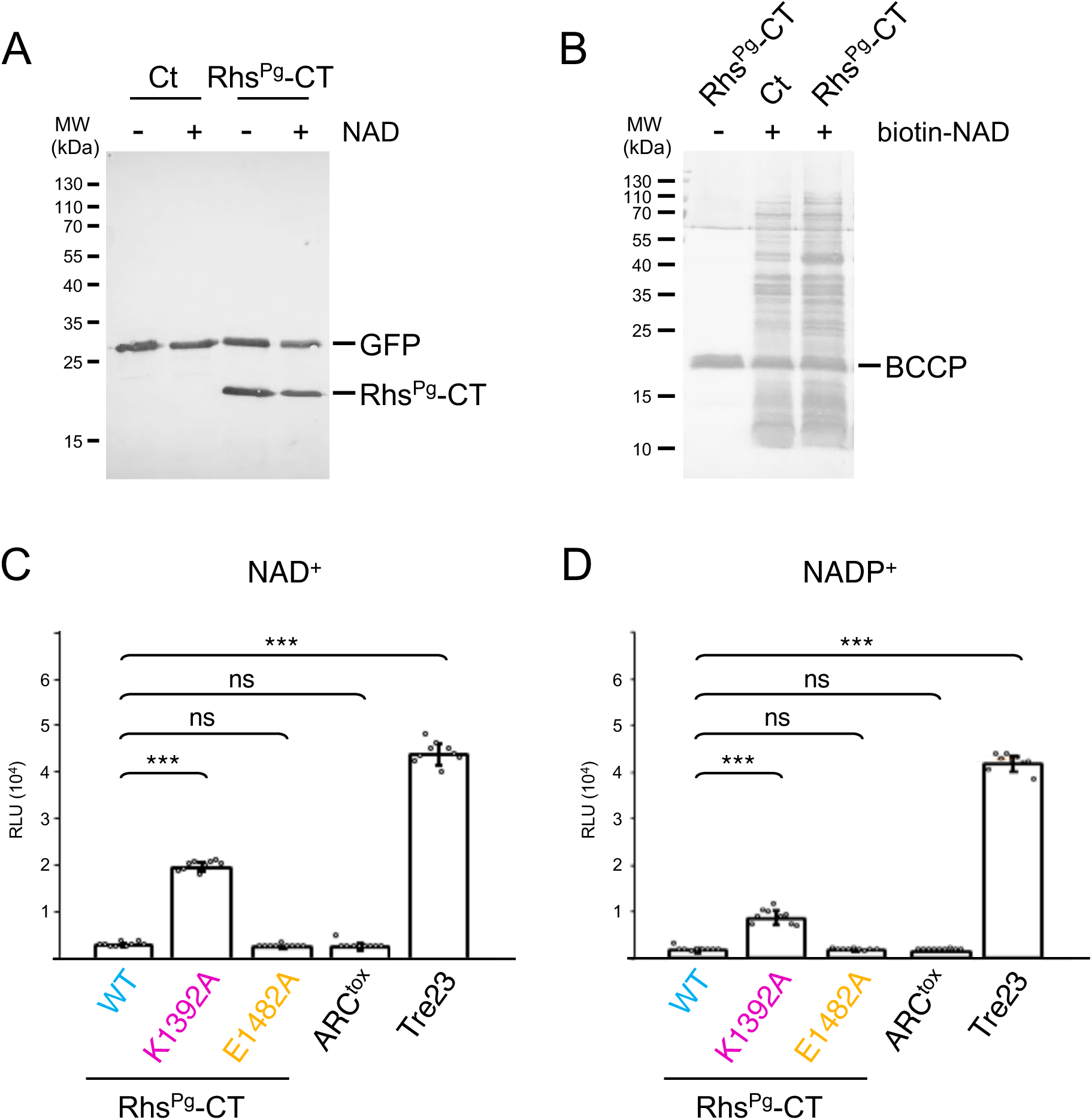
Rhs^Pg^ has no measurable effect on translation or protein ADP-ribosylation but displays NAD(P)⁺ glycohydrolase activity *in vitro*. **(A)** Coupled *in vitro* transcription-translation. *In vitro* transcription-translation of the Strep-tagged GFP reporter protein in reactions supplemented with NAD^+^ and Strep-tagged Rhs^Pg^-CT, as indicated. After 3 h of incubation, proteins were separated by SDS-PAGE, transferred onto nitrocellulose and immunodetected using anti-Streptag antibody. Molecular weight standards (MW, in kDa) are indicated on the left. (**B**) *Ex vivo* protein ADP-ribosylation assays of cellular extracts of *E. coli* in the presence of Rhs^Pg^-CT and of biotin-labelled NAD^+^ (biotin-NAD), as indicated. Proteins were separated by SDS-PAGE, transferred onto nitrocellulose and biotinylated proteins were detected using Streptavidin-Alkaline Phosphatase conjugate. Molecular weight standards (MW, in kDa) are indicated on the left. The only naturally biotinylated protein of *E. coli* (Biotin carboxyl carrier protein, BCCP) is indicated on the right. **(C** and **D)** NAD(P)⁺ glycohydrolase assay. Fluorescence-based assay measuring NAD^+^ (**C**) and NADP^+^ (**D**) levels after incubation of 0.6 mM of NAD^+^ (**C**) or of NADP^+^ (**D**) with 0.1 μM of wild-type (WT) Rhs^Pg^-CT or its K1392A or E1482A variants for 30 min. Control reactions with 0.1 μM of ARC^tox^ (NAD/NADP-consuming) or of Tre23 (ART) were included. RFU, relative fluorescence units. Bars represent the mean of nine independent reactions (each independent value indicated by circles). Standard deviations are indicated. *** denotes a statistically significant difference between the two conditions, based on an unpaired *t*-test (P<0.001). ns, non-significant.

### Rhs^Pg^-CT has NAD(P)⁺ glycohydrolase activity

The previous data suggested that if Rhs^Pg^-CT engages NAD⁺, it may preferentially catalyze NAD⁺ hydrolysis rather than ADP-ribose transfer to protein or RNA targets. To test this, we quantified NAD/NADP glycohydrolase (NADase) activity directly using a fluorescence-based assay. NAD⁺ or NADP⁺ levels were measured after 30 min of incubation with Rhs^Pg^-CT obtained from IVT. Rhs^Pg^-CT strongly depleted both NAD⁺ and NADP⁺, at levels comparable to the *Pantoea ananatis* ARC ADP-ribosylcyclase (ARC^tox^, 47), whereas the *Photorhabdus laumondii* ART Tre23 toxin had no measurable effect (Fig. 4*C*). The K1392A substituted Rhs^Pg^-CT yielded an intermediate phenotype with partial depletion of the NAD(P)⁺ pool while the E1482A mutation had no impact, mirroring the cell toxicity of these variants. These results indicate that the *P. gergoviae* Rhs C-terminal domain is an ART-like fold effector that functions primarily as a NAD(P)⁺ glycohydrolase.

### Phylogenetic analyses define a new ART-related NADase clade, Tne5

To position Rhs^Pg^-CT within the tree of T6SS-associated NAD(P)⁺-consuming effectors, we conducted phylogenetic analyses. Maximum-likelihood trees built from representative members of the four previously defined Tne families (Tne1-Tne4) as well as characterized NAD^+^-dependent toxins (*Mycobacterium tuberculosis* Tuberculosis Necrotizing Toxin (TNT, 59), *Aspergillus fumigatus* AfNADase (60), *P. ananatis* ARC^tox^ (47) and *P. laumondii* ART Tre23 (38)) revealed that Rhs^Pg^-CT does not cluster with any of these groups but instead defines a distinct clade (Fig. 5*A*). We therefore propose that this lineage represents a new family, which we designate Tne5, and its cognate immunity protein Tni5. Interestingly, the Tne5 clade branches closer to canonical ADP-ribosyltransferases (ARTs) than to other Tne families, including proximity to the *P. lamunondii* Tre23 effector (Fig. 5*A*). This evolutionary relationship is mirrored at the structural level: Tne5 shares the same arrangement of β-strands as ART enzymes, whereas Tne1-Tne4 display an alternative strand topology (Fig. S4).

**Figure 5.**
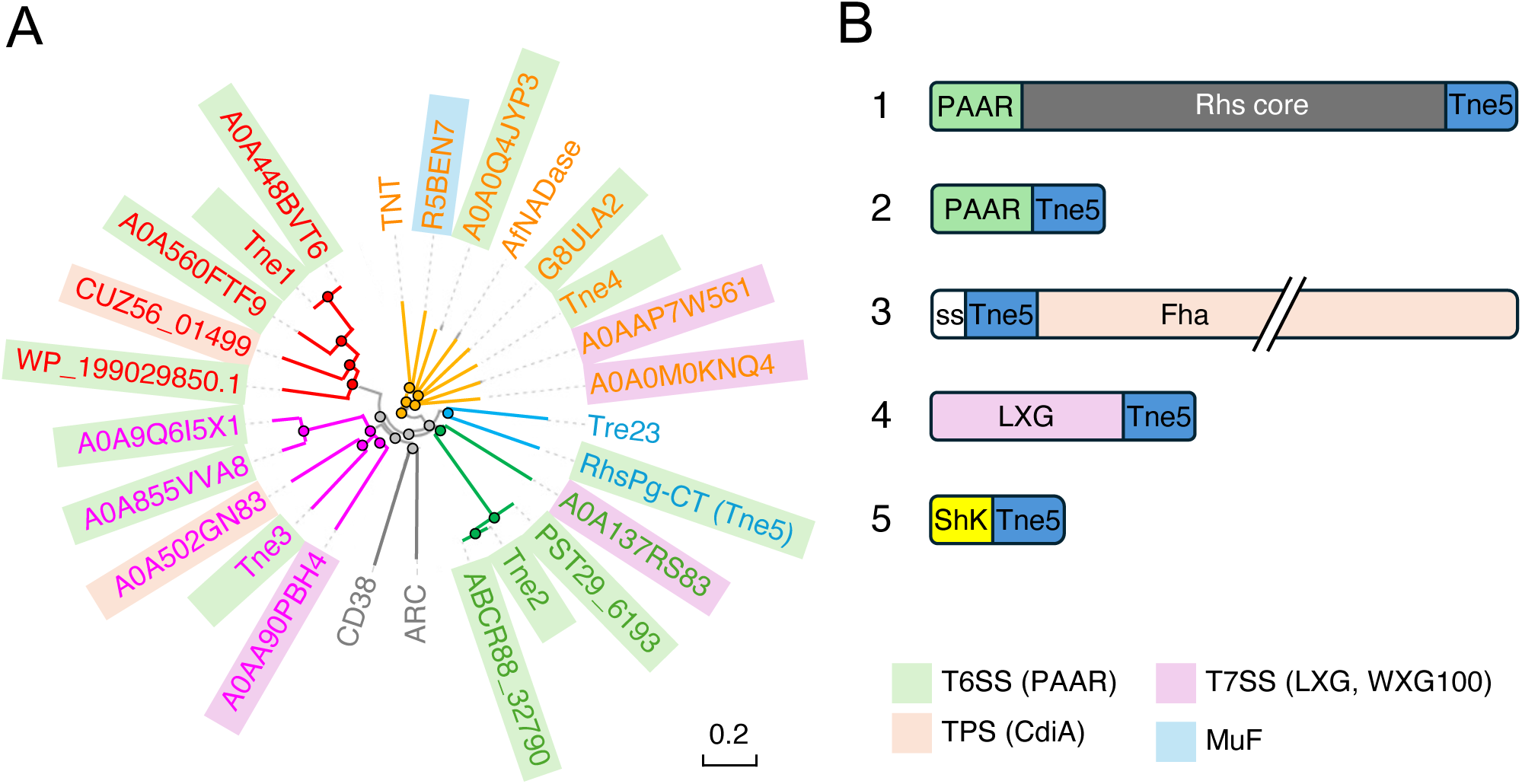
Phylogenetic and genomic analyses define Tne5 as a new family of ART-related NAD(P)⁺ glycohydrolases. (**A**) Maximum-likelihood phylogenetic tree of prototypical and representative members of the Tne1 (red lettering), Tne2 (green lettering), Tne3 (pink lettering) and Tne4 (orange lettering) NADase families, canonical NADases (TNT, AfNADase), ADP-ribosylcyclases (ARC^tox^ and human CD38), Tre23 ADP-ribosyltransferase, and Rhs^Pg^-CT. Toxin associated with T6SS, CDI, T7SS and MuF systems are underlined in grey, flesh, pink and blue, respectively. Rhs^Pg^-CT clusters in a distinct clade, separate from the previously defined Tne1-Tne4 families. (**B**) Representative genomic contexts of Tne5 domains identified by JackHMMER searches. Tne5 domains occur in five major architectures, as C-terminal extensions of polymorphic toxins associated with T6SS-dependent PAAR (green) and Rhs (grey) domains, contact-dependent growth inhibition (CDI, flesch), T7SS-dependent LXG domains (pink), or ShK elements (yellow).

To assess the distribution of Tne5-like effectors, we performed iterative homology searches using JackHMMER with the *P. gergoviae* Rhs^Pg^ Tne5 sequence as a query. This analysis identified bacterial Tne5 homologues associate with T6SS PAAR or PAAR-Rhs domains, contact-dependent inhibition (CDI) systems, type VII secretion systems (T7SS), as well as with ShK domains found associated with toxins in nematodes and sea anemones (Fig. 5*B*).

## DISCUSSION

In this study, we demonstrated that *Pluralibacter gergoviae* encodes a T6SS gene cluster, bearing all the core genes, essential for the assembly of a functional T6SS (Fig. 1*A*). This T6SS is active in laboratory conditions and displays antibacterial activity against *E. coli* cells (Fig. 1*B*). Genomic analyses suggest that this T6SS gene cluster carries two toxin-immunity pairs, including a putative Tae4 amidase and its cognate immunity Tai4, and a Rhs polymorphic toxin followed by a small open reading frame encoding a cytosolic protein. (Fig. 1*A*) This Rhs protein is predicted to comprise a PAAR domain followed by a bundle of transmembrane helices, a Rhs core and a C-terminal extension (Fig. 1, *C* and *D*). This organization is typical of Rhs polymorphic toxins delivered by the T6SS, in which the PAAR domain specifies association to the VgrG spike protein (56–58). Indeed, AlphaFold3 modeling suggests formation of a VgrG-Rhs complex involving binding of the PAAR domain to the VgrG tip (Fig. 1*E* and Fig. S2). In addition, transmembrane helices are usually protected by chaperones of the EagR family (21, 22), and such a putative chaperone is encoded upstream the *rhs* gene (Fig. 1, *A* and *E*, Fig. 2). Consistent with this architecture, heterologous expression of the Rhs C-terminal extension in *E. coli* was toxic and co-expression of the downstream gene restored viability (Fig. 2), demonstrating that *rhs* and the downstream gene encode a *bona fide* polymorphic toxin-immunity pair.

The AlphaFold3 model of the toxic C-terminal Tne5 extension revealed a ADP-ribosyltransferase (ART)-like α/β fold, consisting of a split β-sheet composed of five strands in the order β5-β2 / β1-β3-β6, buttressed by helices to create a cleft in which the NAD/NADP binds (Fig. 3, *A*, *B* and *C*). In this model, the strand usually at position β4 is missing, or more likely not well predicted as an unstructured sequence is present at position of the typical β4. Within this architecture, Lys1392 and Glu1482 are predicted to contribute a basic-acidic pocket that accommodates the NAD molecule in the ligand-bound AlphaFold3 prediction and that may participate to NAD engagement and catalysis (Fig. 3, *B* and *C*). The importance of this pocket is echoed by the Tne5-Tni5 model, which places the immunity protein over the cleft in a way that would sterically occlude access of the NAD (Fig. 3*D*). Mutational analysis however showed that substituting Glu1482 with alanine had only a mild impact on toxicity, whereas the K1392A variant was attenuated (Fig. 3*E*). These results suggest that Lys1392 makes a primary contribution, likely by electrostatic engagement of the pyrophosphate/ribose region or by stabilizing a reaction intermediate, while Glu1482 is not critical for catalytic output. The non-essential role of the canonical acidic residue and the modest effects of the lysine residue mutation indicate that the predicted catalytic pocket does not correspond to a *bona fide* ADP-ribosyltransferase active site. This apparent mismatch between fold-based predictions and catalytic requirements is however in agreement with the functional assays. Although the Tne5 domain adopts an ART-like architecture, it fails to catalyze protein ADP-ribosylation (Fig. 4, *A* and *B*) but instead efficiently depletes NAD⁺ and NADP⁺ *in vitro* (Fig. 4, *C* and *D*). These observations suggest that Tne5 does not operate as a classical ART but rather as an NAD-consuming enzyme, such as an NADase, or as a highly divergent ART in which water, rather than a protein or RNA substrate, acts as the ADP-ribose acceptor. In this context, the predicted Lys1392-Glu1482 pocket may primarily contribute to NAD/NADP binding or positioning without defining a canonical transferase active center. Furthermore, the comparable hydrolysis activity of NAD⁺ and NADP⁺ suggests relaxed dinucleotide specificity of the active site, which is compatible with the geometry of the modeled pocket and with the modest contribution inferred for Glu1482. Further mutagenesis studies would be required to better describe the Tne5 catalytic residues.

The identification of Tne5 as a distinct clade substantially extends the known diversity of T6SS- associated NAD(P)⁺-consuming effectors (Fig. 5*A*). Previous classifications grouped Tne1-Tne4 as NADases that adopt ART-like folds, in which the six β-strands of the split β-sheet are however not organized as in typical ARTs, making them evolutionarily distant from *bona fide* ADP-ribosyltransferases (Fig. S4). In contrast, Tne5 phylogenetic proximity to ART enzymes and its conservation of the ART-like β-sheet topology suggest that Tne5 may represent a more ancestral or transitional lineage between classical ADP-ribosyltransferases and specialized NAD-depleting toxins. This reinforces the notion that ART-like scaffolds provide a versatile platform that can be repurposed toward fundamentally different catalytic outcomes, ranging from macromolecule modification to cofactor hydrolysis.

Taken together, these results demonstrate that Tne5 represents a new addition to the growing repertoire of NAD-targeting enzymes, which are highly potent trans-kingdom toxins as NAD(P)⁺ cofactors play an essential and conserved role in the metabolism of all living organisms. Tne5-like domains are broadly distributed within T6SS, CDI, and T7SS but also fused to ShK modules found in eukaryotic toxins from nematodes and sea anemones, (Fig. 5*B*), highlighting their modularity and evolutionary success as polymorphic toxins in a wide range of biological and ecological contexts. This mirrors recent observations for ADP-ribosyl cyclases (46, 47) and supports a unifying view in which NAD(P)⁺ depletion constitutes a central antibacterial strategy recurrently selected for delivery by distinct secretion platforms.

## EXPERIMENTAL PROCEDURES

### Bacterial strains, growth conditions, media and chemicals

Strains used in this study are listed in Table S1. *Pluralibacter gergoviae* ATCC33028 was kindly provided by A. Davin-Régli (MCT, Marseille, France)*. E. coli* strains DH5α and CC118λpir were used for cloning, whereas *E. coli* W3110 was used for toxicity and bacterial competition assays. *P. gergoviae* and *E. coli* cells were grown at 37°C in Lysogeny Broth (LB) with agitation or on LB agar (1.5%) plates. When needed, media were supplemented with ampicillin (100 μg.mL^-1^), chloramphenicol (30 μg.mL^-1^), or streptomycin (100 μg.mL^-1^). Gene expression was induced by the addition of 0.2% of L-arabinose or of 0.05 mM of IPTG, or repressed with 1% of glucose.

### Plasmid construction

Plasmids and oligonucleotides used in this study are listed in Table S1. All cloning procedures were performed using Sequence and Ligation Independent Cloning (SLIC). Oligonucleotides were purchased from IDT. DNA fragments encompassing *tssL* or encoding Rhs^Pg^-CT or the Tni5 immunity were amplified from *P. gergoviae* genomic DNA using the Q5 polymerase (NEB). Vector backbones and full plasmid templates were amplified using the Q5 polymerase (NEB). PCR products were purified using NucleoSpin Gel and PCR Clean-up columns (Macherey-Nagel), digested with the appropriate restriction enzymes (NEB), and repurified prior to ligation overnight at 16°C with T4 DNA ligase (NEB). Ligated plasmids were transformed into *E. coli* DH5α of CC118λpir and screened by colony PCR using EconoTaq PLUS GREEN 2× Master Mix (Biosearch Technologies). Plasmid DNA was extracted using the NucleoSpin Plasmid kit (Macherey-Nagel) and verified by Sanger sequencing (Eurofins). Constructs were selected on LB agar plates containing ampicillin (for pNDM220), chloramphenicol (for pBAD33), or streptomycin (for pKNG101). Glucose (1%) was added for selection with the pNDM220 vector. Site-directed mutagenesis of *rhs^Pg^-CT* was performed using SLIC primers carrying the desired codon mutations for a PCR reaction amplifying the whole plasmid. After treatment with T4 DNA polymerase, plasmids were annealed and transformed into DH5α competent cells. All plasmids were checked by colony-PCR and verified by DNA sequencing (Eurofins).

### Construction of the isogenic tssL mutant of P. gergoviae

The *P. gergoviae* isogenic *tssL* mutant strain, consisting at the insertion of two stop codons at codons 2 and 3, was constructed as follows. Two 600-bp fragments corresponding to the sequences upstream and downstream of the codons to be mutated were generated by PCR using *P. gergoviae* chromosomal DNA as template, the Q5 DNA polymerase and primers pKNG-TssL-SLIC1 and TssL-STOPSTOP-SLIC-2, and TssL-STOPSTOP-SLIC-3 and TssL-pKNG-SLIC-4 and. These two PCR products overlap on a 20-bp region and are centered on the mutated codons. Both fragments were cloned into the suicide vector pKNG101 digested with BamHI using SLIC using the T4 DNA polymerase, yielding pKNG101-tssLSTOP. pKNG101- tssLSTOP was introduced into *P. gergoviae* by conjugation using the MFDpir conjugative strain. The first event of homologous recombination was selected on LB-agar plate supplemented with streptomycin. Several transconjugants were streaked onto LB agar without NaCl supplemented with 6 % of saccharose to select the second recombination event. After incubation for 48 h at 12°C, plates were incubated ON at 37°C. Several clones were then tested for streptomycin sensitivity and checked by colony-PCR.

### Interbacterial competition assay

Antibacterial activity was measured using the chlorophenol-red β-D-galactopyranoside (CPRG)-based lysis-associated β-galactosidase assay (LAGA) as previously described (61). Briefly, this assay is based on the degradation of chlorophenol-red β-D-galactopyranoside (CPRG), a membrane-impermeable chromogenic substrate of the β-galactosidase, which is cleaved by the β-galactosidase released by the lysis of *E. coli* W3110 *lacZ^+^* recipient cells. Attacker *P. gergoviae* (WT or Δ*tssL*) and recipient *E. coli* W3110 strains were grown separately to an OD_600_ of 0.5 in LB medium supplemented with 0.05 mM IPTG, normalized to identical optical densities, and mixed to a 1:4 (donor:recipient) ratio. Ten µL of the mixtures were spotted onto LB agar plates supplemented with 0.05 mM of IPTG. Plates were incubated for 4 hours at 37 °C, and recipient cell lysis was observed by the addition of a 10-µL drop of 2 mM of CPRG (Roche) on the top of each spot.

### Toxicity assay in E. coli

*E. coli* DH5α cells were co-transformed with the pNDM220 and pBAD33 plasmid pairs bearing either wild-type and mutant Rhs^Pg^-CT, and Rhs^Pg^ immunity, respectively, or empty vectors as controls. Transformants were selected on LB agar plates supplemented with ampicillin, chloramphenicol, and 1 % of glucose. Overnight cultures grown in the presence of antibiotics and glucose were diluted in 10-fold series, and 10-μL drops were spotted on LB agar plates containing antibiotics and either 1% glucose (repression conditions) or 0.05 mM IPTG (toxin induction from pNDM220) and 0.2% L-arabinose (immunity induction from pBAD33). Plates were incubated at 37 °C for 16-20 hours before imaging.

### In vitro coupled transcription-translation assay

Coupled *in vitro* transcription-translation assays were performed as previously described (38, 39) using the PURExpress® In vitro Protein synthesis kit (NEB). Tne5 was obtained using a DNA template amplified using the 5’UTR-Tne5Pg and 3’UTR-Tne5Pg-strep primers. A second coupled *in vitro* transcription-translation reaction was then performed with a DNA template coding for the GFP-strep reporter protein, amplified using the 5’UTR-GFP and 3’UTR-GFP-strep primers. When indicated, reactions contained 0.1 mM NAD^+^ and/or 1 μL of IVT-produced Tne5. Reactions were performed for 2 hours at 37 °C, proteins were separated by SDS-PAGE, transferred onto nitrocellulose membranes, and GFP-strep and Tne5-strep were detected by immunoblotting with strep-Tag antibodies (Classic, BioRad).

### Ex-vivo biotin-NAD protein labeling assay

Fifty mL of *E. coli* DH5α cells were grown to late exponential phase and resuspended in 0.5 mL of Phosphate-buffered saline (PBS) buffer. Cells were then disrupted by sonication and unbroken cells were discarded. 0.1 mM of 6-biotin-17-NAD^+^ and 1 μL of IVT-produced Tne5 were added, when indicated, to 30 μL of cell lysate. After 30 min of incubation, an equal volume of 2× Laemmli loading dye was added, heated for 5 min at 95 °C, prior to protein separation by SDS-PAGE, transfer onto nitrocellulose membranes and immunodetection of biotin-ADP-ribosylated proteins with streptavidin-alkaline phosphatase conjugate (Invitrogen).

### In vitro NAD(P)^+^ degradation assay

Reactions were performed in 100 μL of PBS supplemented with 0.6 mM of NAD^+^ or of NADP^+^ (Roche). One μL of wild-type and mutant Tne5 domains obtained from the *in vitro* coupled transcription-translation reaction were added. Samples were incubated at room temperature for 30 min. The reaction was then quenched by adding 50 μL of 6 M NaOH. Fluorescence was measured using a TECAN plate reader (excitation: 340 nm; emission: 420 nm) after 30 min of incubation at room temperature in the dark. Controls including the NAD-consuming ARC^tox^ domain (47) and the Tre23 ART domain (38). NAD/NADP hydrolysis data were analyzed using an unpaired *t*-test. All analyses were performed using the GraphPad Prism software.

### Bioinformatics analyses and AlphaFold3 modeling

The T6SS locus was identified in the ATCC 33028 genome assembly (GenBank accession: GCA_001598855.1) by BLASTP against canonical T6SS components. Domain architecture of the Rhs protein was annotated with HHpred and PFAM; transmembrane helices were predicted with TMHMM. Phylogeny reconstruction of the Tne tree was performed using the Simple Phylogeny server (EMBL-EBI, 62) based on a multiple sequence alignment provided by Clustal Ω 1.2.4 (EMBL-EBI) using selected members of the Tne1-4 families and previously characterized NADases (*Mycobacterium tuberculosis* TNT (59), AfNADase (60)), ADP-ribosyl cyclase (ARC^tox^, 47) and the Tre23 ART (38). Tne5 closest homologues were retrieved using JackHMMER (EMBL-EBI, 63) using a single iteration. AlphaFold3 model predictions were run on the AlphaFold server using default MSA generation (64). The toxin-immunity complex was predicted with AF3 multimer mode (stoichiometry 1:1). Models were visualized in ChimeraX. pLDDT and PAE as well as confidence metrics are indicated in the corresponding figures.

## Supporting information

This article contains supporting information (four Supplemental Figures and one Supplemental Table).

## Acknowledgments

We thank Anne Davin-Régli for providing strain *Pluralibacter gergoviae* ATCC33028, Dukas Jurėnas for discussions regarding the ART/Tne fold and catalytic residues, members of the Cascales laboratory, Anne Galinier and Yohann Duverger and the JBD thesis committee for discussions and support, and Melissa Guilbert and Audrey Gozzi for technical support.

## Author contributions

J. B. D., and E. C., conceptualization; J. B. D., and E. C., methodology; J. B. D., and E. C., validation; J. B. D., formal analysis; J. B. D., M. D., and E. C., investigation; J. B. D., and M. D., writing—review & editing; J. B. D., and E. C., visualization; E. C., supervision; J. B. D., and E. C., resources, E. C., project administration; E. C., writing—original draft; J. B. D., and E. C., funding acquisition.

## Funding and additional information

This work was funded by the Centre National de la Recherche Scientifique, the Aix-Marseille Université, and by a grant from the Agence Nationale de la Recherche (ANR-24-CE44-7545) and the Fondation Bettencourt-Schueller. JBD was supported by a doctoral fellowship and a 4^th^ year extension from the Fondation pour la Recherche Médicale (contracts ECO202206015565 and FDT202504020886).

## Abbreviations

The abbreviations used are:

ARC: ADP-ribosyl cyclase
ART: ADP-ribosyl transferase
CDI: contact-dependent growth inhibition
Hcp: haemolysing coregulated protein
NAD: nicotinamide adenine dinucleotide
NADase: NAD(P)⁺ glycohydrolase
NADP: nicotinamide adenine dinucleotide phosphate
PAAR: Proline-Alanine-Alanine-Arginine
PT: polymorphic toxin
Rhs: Rearrangement hot spot
T6SS: type VI secretion system
T7SS: type VI secretion system
Tag: type VI secretion-associated gene
Tne: type VI NADase effector
Tni: type VI NADase immunity Tss type VI secretion
VgrG: Valine-glycine repeat G.

## LEGEND TO SUPPLEMENTAL FIGURES

**Supplemental Figure 1. AlphaFold3 structural model of full-length Rhs^Pg^.** *A,* Structural model of Rhs^Pg^ colored by predicted Local Distance Difference Test (pLDDT) scores, ranging from blue (high confidence, >90) to orange (low confidence, <50). *B,* Predicted Aligned Error (PAE) plot of Rhs^Pg^ showing the expected position error (in Å) for each residue pair (dark green, high confidence; light green, low confidence).

**Supplemental Figure 2. Predicted assembly of the Rhs^Pg^ effector with VgrG and the cognate EagR-like chaperone.** *A,* AlphaFold3 multimer model of the effector-loaded spike complex including a trimer of VgrG (shades of orange), a dimer of EagR chaperones (purple), and the Rhs^Pg^ effector. The different Rhs^Pg^ domains are colored as in Fig. 1, *C* and *D* (PAAR domain, light green; TMD, dark green; Rhs core, grey; C-terminal extension, blue). A close-up view of the complex, emphasizing the interface between the VgrG tip and the Rhs^Pg^ PAAR domain, as well as the protection of the TMD by the EagR dimer, is shown in the inset on left. *B,* AlphaFold3 multimer model of the effector-loaded spike complex colored by pLDDT confidence scores. The overall interfacial confidence score (ipTM, 0.47) is indicated. ipTM scores of VgrG-PAAR and EagR-TMD subcomplexes are 0.66 and 0.80, respectively. *C,* Predicted aligned error (PAE) plot of the effector-loaded spike complex.

**Supplemental Figure 3. AlphaFold3 structural models of the Tne5 toxin domain alone and in complex with ligand or immunity.** *A, C, and E,* Predicted structures of the Rhs^Pg^-CT domain alone (*A*), bound to NADH (*C*) and in complex with its cognate immunity protein (*E*) colored by pLDDT. Confidence scores (pTM or piTM) are indicated. *B, D, and F,* PAE plots of the Rhs^Pg^-CT domain alone (*B*), bound to NADH (*D*) and in complex with its cognate immunity protein (*F*).

**Supplemental Figure 4. β-sheet topology distinguishes Tne5 from other Tne families.** *A and B,* Topological diagrams of the conserved core β-sheets of Tne1-Tne4 (*A*) and Tne5 and ARTs (*B*). The diagrams represent the conserved catalytic cores and do not include additional peripheral structural elements. β-strands are rainbow-colored, from blue (β1) to red (β6). Whereas all Tne families share the 6-stranded β-sheet, the organization of the β-strands differ significantly between the Tne1-4 families and Tne5, which shares the same arrangement as ARTs. N, N-terminus; C, C-terminus.

