## Supplemental Information for "An ART-fold Rhs toxin from *Pluralibacter gergoviae* defines Tne5, a new clade of NAD(P)⁺ glycohydrolases effectors"

### Table S1. Strains, plasmids and oligonucleotides used in this study.

| STRAINS | | |
|---|---|---|
| **Strains** | **Description** | **Source/Reference** |
| *Pluralibacter gergoviae* | | |
| *P. gergoviae* ATCC 33028 | Wild-type *Pluralibacter gergoviae* NBRC 105706 (ATCC 33028) | Brenner *et al.*, 1980 |
| Δ*tssL* | insertion of two consecutive STOP codons after the START codon of *tssL* (A8H26_06215) | This study |
| *E. coli* K-12 | | |
| DH5α | $F^-$, Δ*(argF-lacZ)*U169, *phoA, supE44,* Δ(lacZ)M15, *relA, endA, thi, hsdR* | New England Biolabs |
| CC118λpir | Δ*(ara-leu) araD* Δ*lacX galE galK phoA thi rpsE rpoB argE recA λpir* phage lysogen | Herrero *et al.*, 1990 |
| W3110 | $F^-$, $\lambda^-$ *rph*1 INV(*rrnD*, *rrnE*) | Laboratory collection |
| MFDpir | *MG1655* RP4-2-Tc*::*[ΔMu1*::aac(3)IV* Δ*aphA* Δ*nic35* Δ*Mu2::zeo* Δ*dapA::(erm-pir)* Δ*recA* | Ferrières *et al.*, 2010 |

| PLASMIDS | | |
|---|---|---|
| pKNG101 | R6K origin, TRK2 origin, mobRK2, *sacB*$^+$, $Sm^R$ | Kaniga *et al.*,1991 |
| pKNG101-tssLSTOP | pKNG101 bearing flanking regions to insert two STOP codons in *tssL* | This study |
| pNDM220 medRBS | Mini-R1, single-copy vector, $LaqI^q$ , $P_{A1/04/03}$, with an attenuated RBS, $Amp^R$ | Laboratory collection |
| pNDM-Tne5$^{Pg}$ | *P. gergoviae tne5* (Rhs amino acids 1330–1503) cloned into pNDM220 medRBS | This study |
| pNDM-Tne5$^{Pg}$ K1392A | Lys1392 to Ala substitution in pNDM-Tne$_{Pg}$ | This study |
| pNDM-Tne5$^{Pg}$ E1482A | Glu1482 to Ala substitution in pNDM-Tne$_{Pg}$ | This study |
| pBAD33 RBS | Expression vector, AraC, arabinose-inducible, Ribosome binding-site, $Cm^R$ | Laboratory collection |
| pBAD-Tni5$^{Pg}$ | *P. gergoviae tni5* (A8H26_06350) cloned into pBAD33 RBS | This study |

| OLIGONUCLEOTIDES | |
|---|---|
| Construction of mutation in *P. gergoviae* [a] | |
| pKNG-TssL-SLIC-1 | CCCTGCAGGTCGACGGATCCCGAATTTACCCTGCTGACGC |
| TssL-STOPSTOP-SLIC-2 | AGCCTGCTGTTGCTGCATAG |

| | |
|---|---|
| TssL-STOPSTOP-SLIC-3 | CTATGCAGCAACAGCAGGCTTAGTAAAGCACGCAGGAACAACAGGC |
| TssL-pKNG-SLIC-4 | CTTATGGTACCCGGGGATCCTGGTAGATTTTCGCCAGCACC |
| Verif-TssL-STOPSTOP | AGCAACAGCAGGCTTAGTAA |
| Plasmid construction [b,c] | |
| 5-pNDMmedrbs-SpeI-Tne5$^{Pg}$ | GATCACTAGT**ATG**AGTTGTAAGAATAGTTGGAATGAATTTCAGAGTAG |
| 3-pNDMmedrbs-EcoRI-Tne5$^{Pg}$ | GATCGAATTCTCATATATTACTATTTGGCAATTTAGATATCACTGCCTCC |
| 5-pBAD33-SalI-Tni5$^{Pg}$ | GATCGTCGACATGAAAGAGGTTATATTTAGTAAGTACGGTATAGATATTC |
| 3-pBAD33-HindIII-Tni5$^{Pg}$ | GATCAAGCTTTTATTTAATATAAGGCTTATATCCATTATCTCTATTCTGAGACGC |
| Site-directed mutagenesis [d] | |
| Tne5$_{Pg}$-K1392A-SLIC1 | AGATACTCCACCATCAAATTGCGC |
| Tne5$_{Pg}$-K1392A-SLIC2 | TTGATGGTGGAGTATCTGCGATTGCATGGGGAGTTAATCGTGC |
| Tne5$_{Pg}$-E1482A-SLIC1 | CGTCACCGGTGTTTGGTCAATAAC |
| Tne5$_{Pg}$-E1482A-SLIC2 | ACCAAACACCGGTGACGGCGGCAGTGATATCTAAATTGCCAAATAG |
| DNA fragments for *in vitro* transcription-translation [e] | |
| 5'UTR-GFP | GCGAATTAATACGACTCACTATAGGGCTTAAGTATAAGGAGGAAAAAATATGAGTAAAGGAGAAGAACTTTTCAC |
| 3'UTR-GFP-strep | AAACCCCTCCGTTTAGAGAGGGGTTATGCTAGTTATTATTTTTCGAACTGCGGGTGGCTCCATTTGTATAGTTCATCCATGCCA |
| 5'UTR-Tne5$^{Pg}$ | GCGAATTAATACGACTCACTATAGGGCTTAAGTATAAGGAGGAAAAAATATGAGTTGTAAGAATAGTTGGAATGAATTTCAG |
| 3'UTR-Tne5$^{Pg}$-strep | AAACCCCTCCGTTTAGAGAGGGGTTATGCTAGTTATTATTTTTCGAACTGCGGGTGGCTCCATATATTACTATTTGGCAATTTAGATATCAC |

[a] sequence complementary to the plasmid (SLIC) underlined.
[b] restriction site underlined.
[c] ATG sequence added in bold.
[d] mutagenized codon underlined.
[e] Streptag sequence underlined

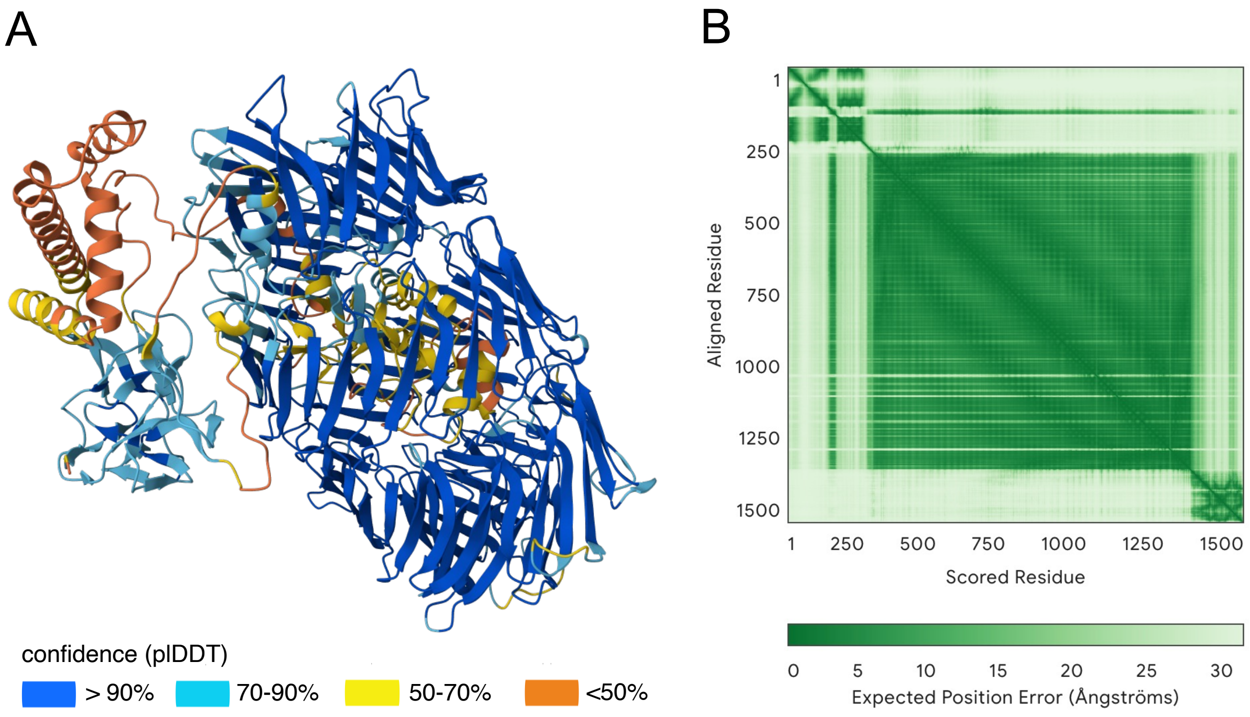


**Supplemental Figure 1. AlphaFold3 structural model of full-length Rhs$^{Pg}$.** (**A**) Structural model of Rhs$^{Pg}$ colored by predicted Local Distance Difference Test (pLDDT) scores, ranging from blue (high confidence, >90) to orange (low confidence, <50). (**B**) Predicted Aligned Error (PAE) plot of Rhs$^{Pg}$ showing the expected position error (in Å) for each residue pair (dark green, high confidence; light green, low confidence).

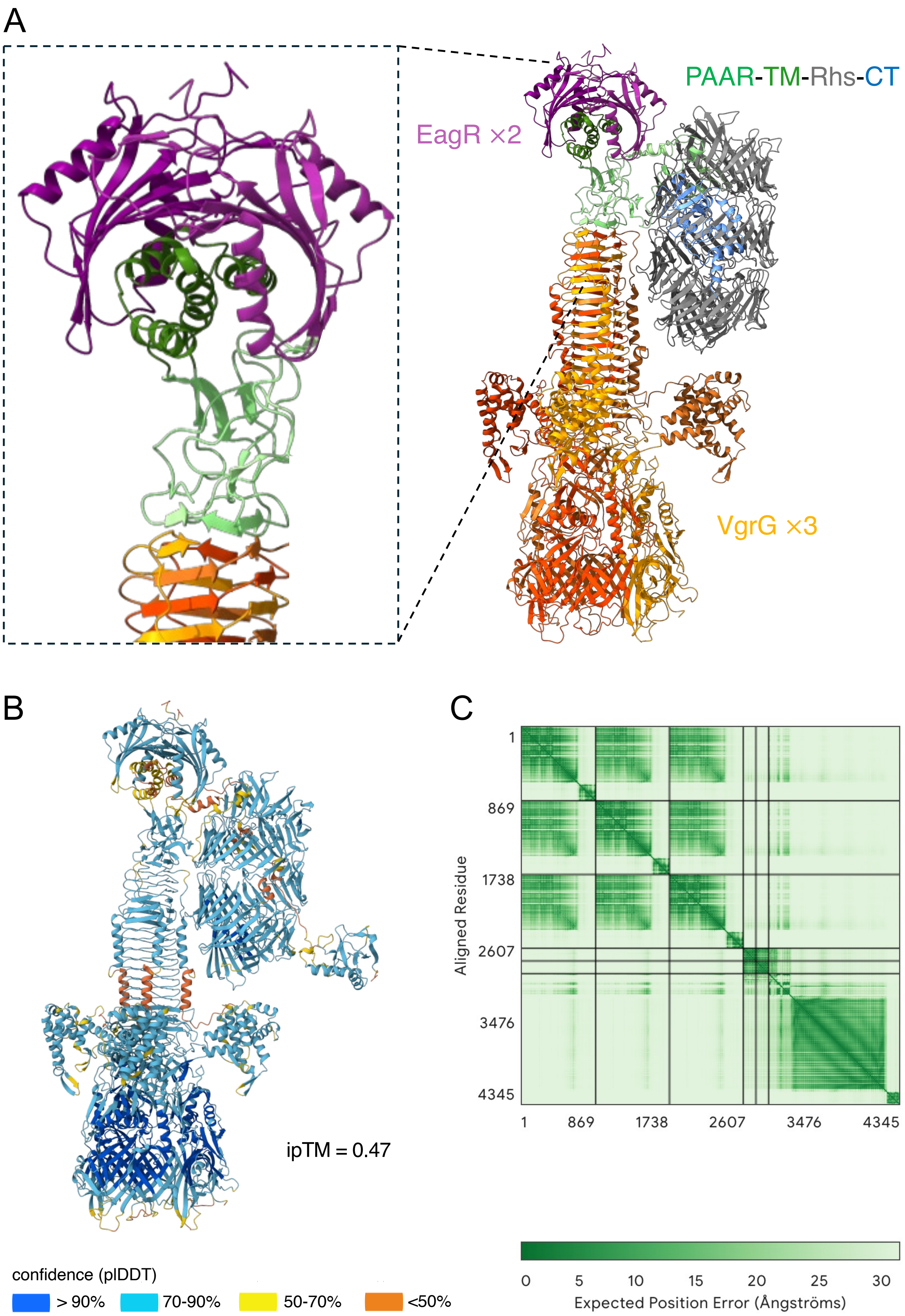
A
PAAR-TM-Rhs-CT
EagR ×2
VgrG ×3
B
ipTM = 0.47
confidence (plDDT)
> 90%
70-90%
50-70%
<50%
C
Aligned Residue
1
869
1738
2607
3476
4345
1
869
1738
2607
3476
4345
0
5
10
15
20
25
30
Expected Position Error (Ångströms)

**Supplemental Figure 2. Predicted assembly of the Rhs$^{Pg}$ effector with VgrG and the cognate EagR-like chaperone.** (**A**) AlphaFold3 multimer model of the effector-loaded spike complex including a trimer of VgrG (shades of orange), a dimer of EagR chaperones (purple), and the Rhs$^{Pg}$ effector. The different Rhs$^{Pg}$ domains are colored as in Fig. 1C-D (PAAR domain, light green; TMD, dark green; Rhs core, grey; C-terminal extension, blue). A close-up view of the complex, emphasizing the interface between the VgrG tip and the Rhs$^{Pg}$ PAAR domain, as well as the protection of the TMD by the EagR dimer, is shown in the inset on left. **(B)** AlphaFold3 multimer model of the effector-loaded spike complex colored by pLDDT confidence scores. The overall interfacial confidence score (ipTM, 0.47) is indicated. ipTM scores of VgrG-PAAR and EagR-TMD subcomplexes are 0.66 and 0.8, respectively. **(C)** Predicted aligned error (PAE) plot of the effector-loaded spike complex.

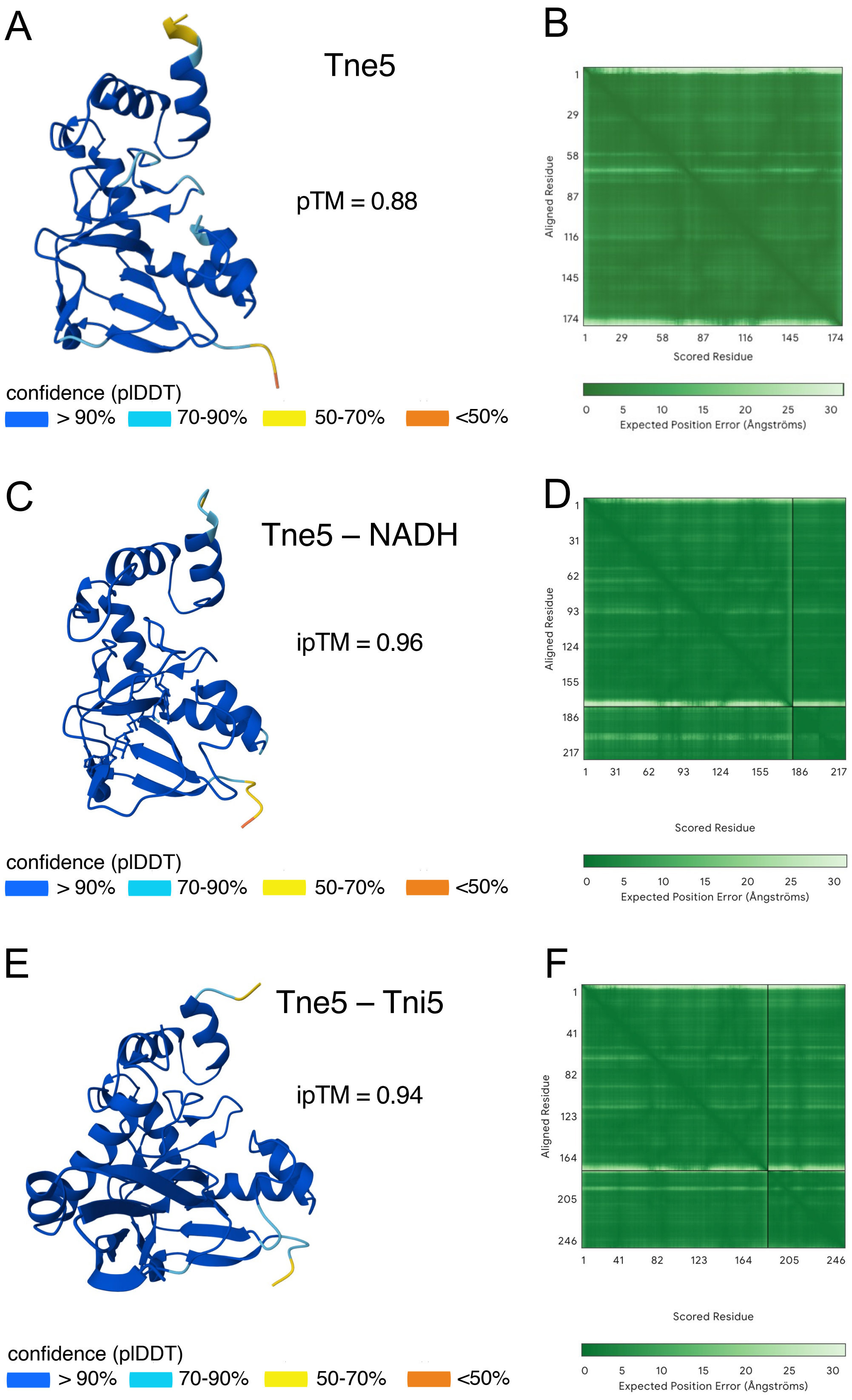


**Supplemental Figure 3. AlphaFold3 structural models of the Tne5 toxin domain alone and in complex with ligand or immunity. (A, C, E)** Predicted structures of the Rhs$^{Pg}$-CT domain alone (**A**), bound to NADH (**C**) and in complex with its cognate immunity protein (**E**) colored by pLDDT. Confidence scores (pTM or piTM) are indicated. (**B, D, F**) PAE plots of the Rhs$^{Pg}$-CT domain alone (**B**), bound to NADH (**D**) and in complex with its cognate immunity protein (**F**).

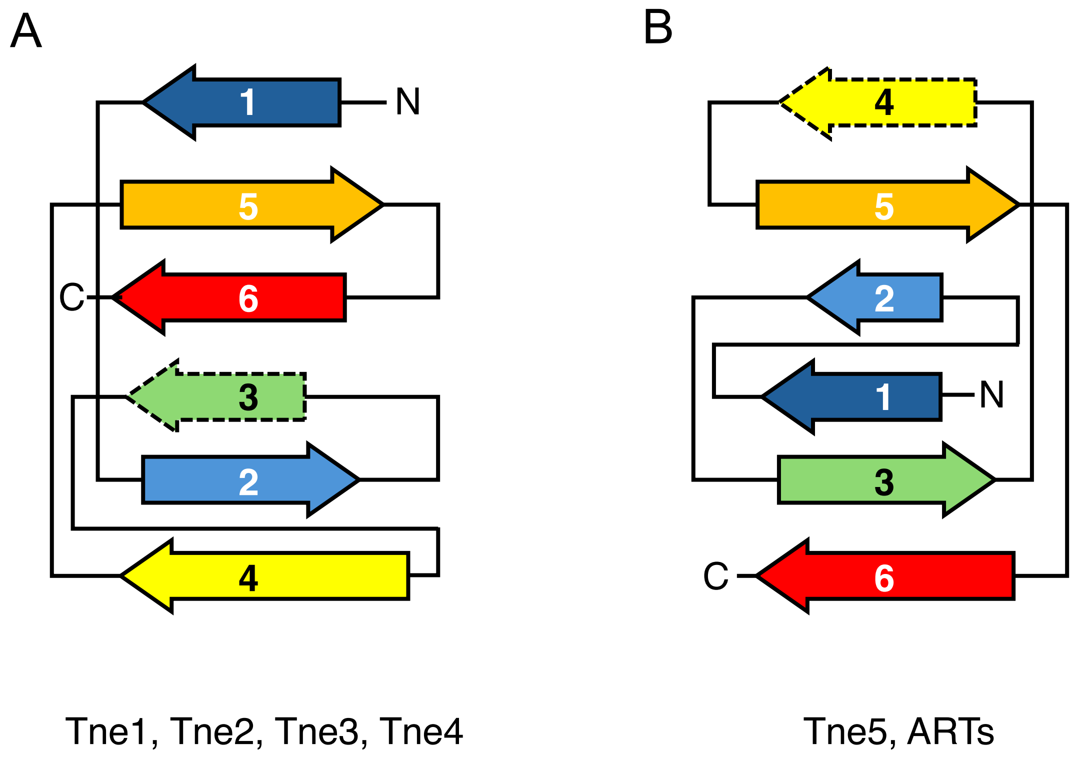


**Supplemental Figure 4. β-sheet topology distinguishes Tne5 from other Tne families.** Topological diagrams of the conserved core β-sheets of Tne1-Tne4 (**A**) and Tne5 and ARTs (**B**). The diagrams represent the conserved catalytic cores and do not include additional peripheral structural elements. β-strands are rainbow-colored, from blue (β1) to red (β6). Whereas all Tne families share the 6-stranded β-sheet, the organization of the β-strands differ significantly between the Tne1-4 families and Tne5, which shares the same arrangement as ARTs. N, N-terminus; C, C-terminus.
